# PAR-Driven Condensation Maintains Stalled Replication Fork Stability

**DOI:** 10.64898/2026.01.07.698279

**Authors:** Lei Zhang, Zeyu Zhang, Timothy R. O’Leary, Arkadi Shwartz, Guoyun Kao, Patrick R. Griffin, Yong Zhang

## Abstract

Poly(ADP-ribose) (PAR) is a nucleic acid-like heterogeneous polymer in nature. Recently, it was found to engage in liquid-liquid phase separation (LLPS), generating condensates as an emerging class of subcellular structures with pivotal functions in response to stimuli. As a post-translational modification catalyzed by PAR polymerases (PARPs), PAR is known to modulate many key events in cells. However, its involvement in biomolecular condensation remains elusive. Through an imaging-based screening of small molecules with diverse biological activities, we here discovered that PAR undergoes LLPS upon inhibiting proteasome in different types of cells, resulting in co-condensation of PAR with proteasome and ubiquitin chains in nucleus. This unprecedented co-condensation is dependent on PARP2 not PARP1 and requires K6-linked ubiquitylation. PAR is shown for the first time to directly interact with ubiquitin chains. Notably, stalled DNA replication forks arose from proteasome inhibition are co-localized with PAR-proteasome-ubiquitin chain condensates. By attenuating replication and stabilizing stalled replication forks, PAR-proteasome-ubiquitin chain condensates sustain genomic integrity under proteasomal stress. This work demonstrates a self-protective mechanism in stressed cells and provides fundamental understanding of PAR condensation in cell biology.

## Introduction

Poly(ADP-ribose) (PAR) is a natural polymer covalently attached to cellular proteins after translation. Protein poly-ADP-ribosylation (PARylation) is catalyzed by PAR polymerases (PARPs), including PARP1, PARP2, PARP5a, and PARP5b.^1–7^ This enzymatic post-translational modification is characterized by sequential transfer of ADP-ribose units from nicotinamide adenine dinucleotide (NAD^+^) donors to protein substrates, producing elongated and branched PAR polymers.^8–12^ The resulting polymer chains are heterogeneous with varied lengths and branching patterns.^13–18^ Such nucleic acid-like modifications can not only alter structures, surface charges, and interactions with other molecules of the PARylated proteins, but also serve as scaffolds for recruiting PAR-specific proteins and assembling signaling or functional complexes, playing vital roles in many events and processes of cells.^19–27^

Biomolecular condensates are identified as dynamic membrane-less compartments in cells, resulted from liquid-liquid phase separation (LLPS) of numerous macromolecules in response to endogenous and environmental stimuli. A variety of proteins and nucleic acids were found to participate in condensation at different subcellular locations.^28–31^ Through compartmentalization and enrichment of structural and functional biomolecules, condensation facilitates signal transduction and biological activities.^32–35^ Owing to its key involvement in diverse cellular functions, biomolecular condensation has been emerging as an increasingly important phenomenon in cell biology.^36–44^

Recent studies reported PAR-mediated biomolecular condensation related to DNA damage response and stress granule formation,^45–48^ suggesting possible existence of other forms of PAR-associated condensates with unknown functions. To explore new PAR-containing condensates, we performed imaging-based screening for U2OS cells treated with small molecules acting on various targets and pathways. Here we unveiled that PAR undergoes LLPS upon proteasomal inhibition in multiple cell lines, forming co-condensates with proteasome and ubiquitin chains in nucleus. PARP2-catalyzed PARylation is required for PAR-proteasome-ubiquitin chain co-condensation. Notably, PAR shows direct interactions with ubiquitin chains seen in condensates and its condensation requires K6-linked ubiquitin chains. Following proteasomal stress, the stability of stalled DNA replication forks is retained by PAR-proteasome-ubiquitin chain co-condensates. This study revealed the formation of PAR-proteasome-ubiquitin chain co-condensation and its protective role in DNA replication under stressed conditions, generating new knowledge of biomolecular condensation in cells.

## Results

### Proteasomal inhibition leads to PAR condensation

As DNA and RNA were frequently observed in cellular condensates, PAR polymer is likely to engage with biomolecular condensation for modulating processes essential for cells under stresses given its nucleic acid-like structure. To this end, we performed PAR immunostaining and imaging in U2OS cells treated with an in-house library of 60 compounds featuring varied molecular targets and biological activities. Quantitative analysis of cellular PAR signal levels and distribution patterns indicated the formation of PAR foci upon treatment by proteasome inhibitors, including MG132, bortezomib (BTZ), and carfilzomib (CFZ) (Fig. S1a). In comparison with the group treated by dimethyl sulfoxide (DMSO), cells following incubation with MG132, BTZ, and CFZ showed rapid development of multiple PAR foci in nucleus under confocal microscope (Fig. 1a), which are characterized by a diameter range of 2-10 µm (Fig. 1b). Remarkably, this phenomenon was observed in other cell lines, including human A2780, LNCaP, and HEK293T and mouse J774A.1 (Fig. S1b-S1e).

**Figure 1.**
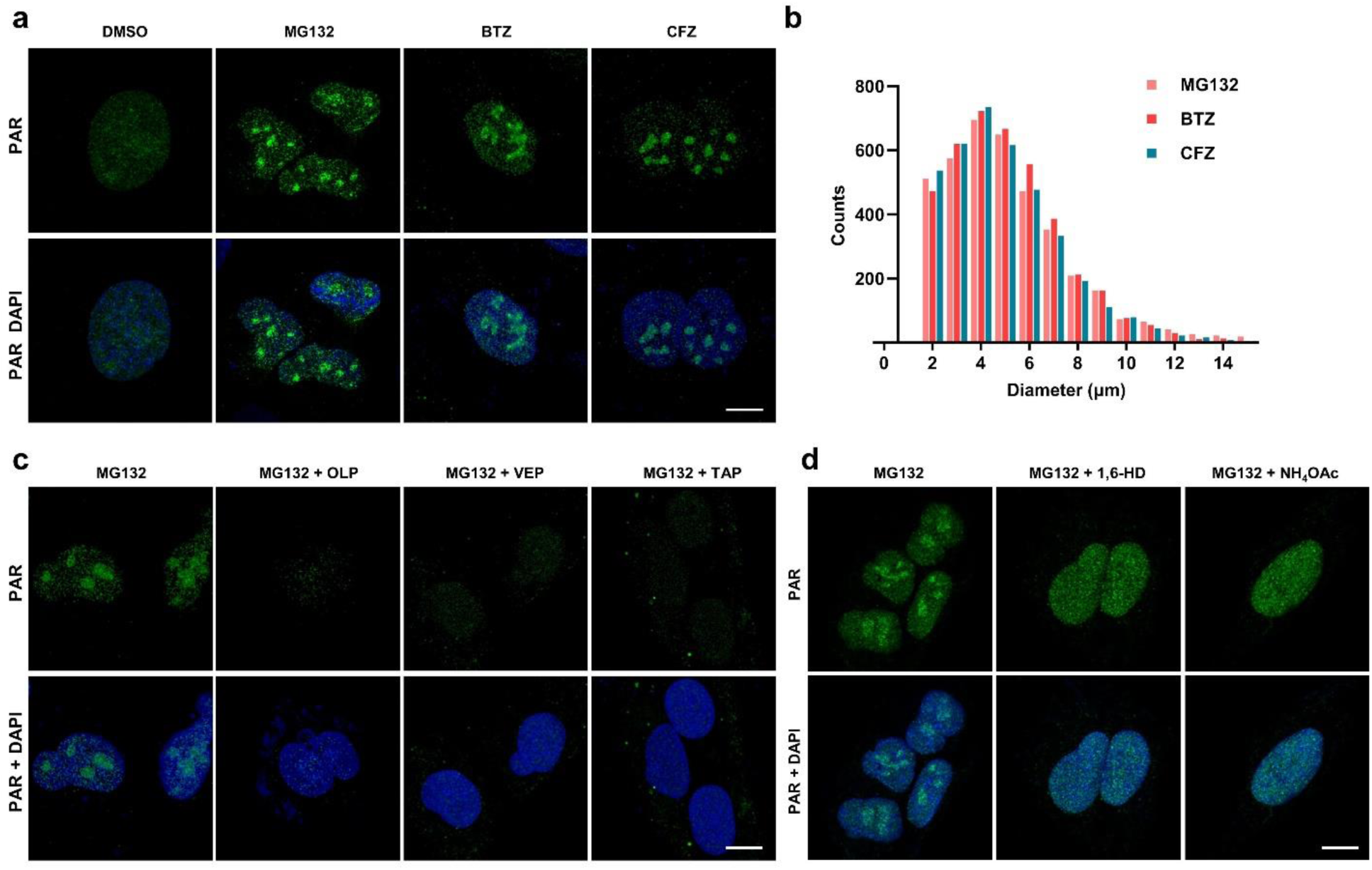
Proteasome inhibitors induce formation of PAR condensates in nucleoplasm. (a) Immunofluorescence imaging of PAR in U2OS cells following treatment for 4 h with DMSO, MG132 (20 μM), bortezomib (BTZ) (10 μM), or carfilzomib (CFZ) (10 μM). (b) The histogram of PAR condensate diameters in U2OS cells after treatment by proteasome inhibitors. A minimum of 5000 PAR foci were analyzed for each condition. (c) Immunofluorescence imaging of PAR condensation in MG132-stimulated U2OS cells without and with pre-treatment by PARP inhibitors. U2OS cells were pre-treated for 0.5 h in the absence and presence of olaparib (OLP) (1 μM), veliparib (VEP) (1 μM), or talazoparib (TAP) (1 μM), followed by 4-h incubation with MG132 (20 μM) and staining for fluorescence imaging. (d) Immunofluorescence imaging of PAR condensates of MG132-treated U2OS cells in the absence and presence of 1,6-hexanediol (1,6-HD) or NH_4_OAc. U2OS cells incubated with MG132 (20 μM, 4 h) were treated without and with 1% 1,6-HD or 0.1 M NH_4_OAc for 0.5 h and then subjected to immunofluorescence staining. Scale bars, 10 μm.

Next, PAR condensation was examined in MG132-treated U2OS cells without and with pre-incubation of PARP inhibitors, including olaparib (OLP), veliparib (VEP), and talazoparib (TAP). Confocal microscopic analysis showed the absence of PAR foci in PARP inhibitor-pretreated cells after blocking proteasomal activity (Fig. 1c), indicating dependency on PARP-catalyzed PARylation for PAR condensation stimulated by proteasome inhibitors. Furthermore, biomolecular condensate-destabilizing reagents 1,6-hexanediol (1,6-HD) and ammonium acetate (NH_4_OAc) were used to validate the generation of PAR foci.^49,50^ Confocal imaging revealed dissolution of PAR foci in MG132-treated cells after additions of 1,6-HD and NH_4_OAc (Fig. 1d). Collectively, these results support cellular PAR condensation in response to proteasomal inhibition.

### Co-condensation of PAR with proteasome

To determine other components present within the observed PAR condensates, we performed PAR pulldown assays for quantitative proteomic studies. Soluble lysates of U2OS cells without and with MG132 treatment were subjected to co-immunoprecipitation using an anti-PAR monoclonal antibody and an IgG isotype control, followed by analysis via liquid chromatography-mass spectrometry (LC-MS) (Fig. 2a, S2a). In contrast to hits identified from DMSO-treated samples, multiple proteasome subunits (e.g. PSMA2, PSMB5, and PSME3) were significantly enriched in samples from MG132-treated cells (Fig. 2b, 2c).

**Figure 2.**
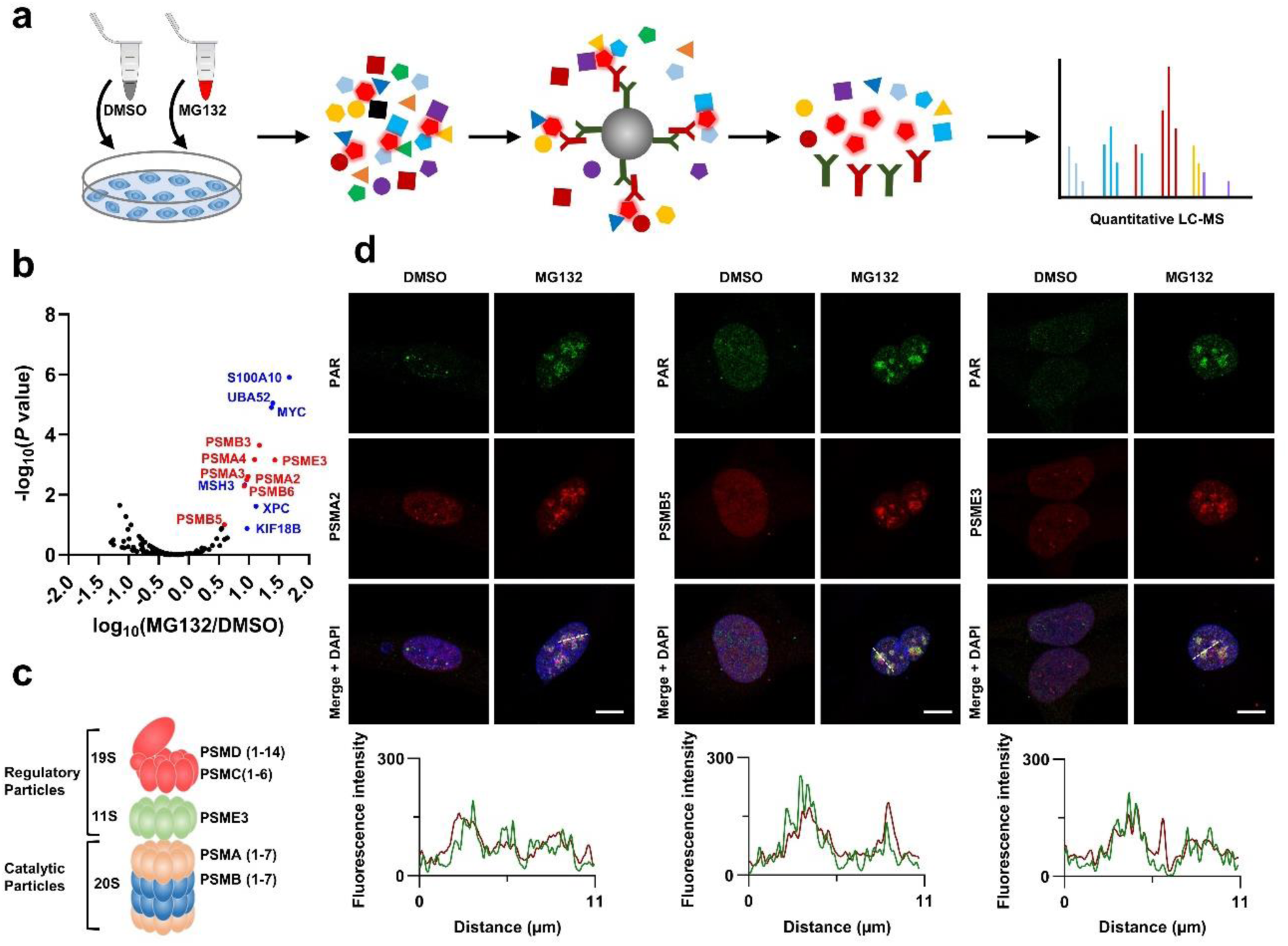
PAR condensates contain proteasome. (a) Schematic of preparation and analysis of PAR pulldown samples by quantitative proteomics. An anti-PAR monoclonal antibody and an IgG isotype control were used for pulldown assays. (b) Volcano plot of PAR condensate-associated proteins in U2OS cells with MG132 treatment. X-axis: protein abundance ratio after stimulation by MG132. Y-axis: *P* value of the protein abundance ratio by t-test. (c) Schematic of 26S proteasome consisting of 19S or 11S regulatory particles and 20S catalytic particles. (d) Confocal microscopy of co-condensates of PAR and proteasome subunits in U2OS cells after 4-h treatment with 20 μM MG132. Plots at the bottom panel represent fluorescence intensities of PAR and respective proteasome subunits from condensates indicated by dashed lines. Scale bars, 10 µm.

Consistent with proteomic results, subsequent confocal microscopy studies indicated the foci development for PSMA2, PSMB5, and PSME3 in MG132-stimulated cells as well as their co-localization with PAR condensates (Fig. 2d). In addition, expression levels of these representative proteasome subunits in U2OS cells were analyzed before and after up to 4-h incubation with proteasome inhibitors. Immunoblot results showed that in comparison with levels of ubiquitin, PAR, and phosphorylated H2AX (Ser139)(p-H2AX) characterized by time-dependent increases post inhibitor exposure, levels of cellular PSMA2, PSMB5, and PSME3 remain largely unchanged over the course of treatment by MG132, BTZ, and CFZ (Fig. S2b). Altogether, these results show that inhibition of proteasome has little effects on its expression levels but triggers co-condensation with PAR polymers.

### PAR-proteasome condensation is dependent on PARP2 not PRAP1

PARP1 and PARP2 with robust auto-PARylation activities account for a majority of cellular PAR production under genotoxic stress.^51,52^ Intriguingly, confocal imaging studies showed weak PARP1 condensation and co-localization with PSMA2 in MG132-treated U2OS cells (Fig. 3a). By contrast, MG132 treatment prompted the formation of PARP2 foci in nucleus and co-condensation with proteasome as imaged by confocal microscopy (Fig. 3b). These findings are consistent with our quantitative proteomic data, in which enrichment of PARP2 but not PARP1 was observed in PAR pulldown samples from MG132-treated cells versus ones from the DMSO-treated group.

**Figure 3.**
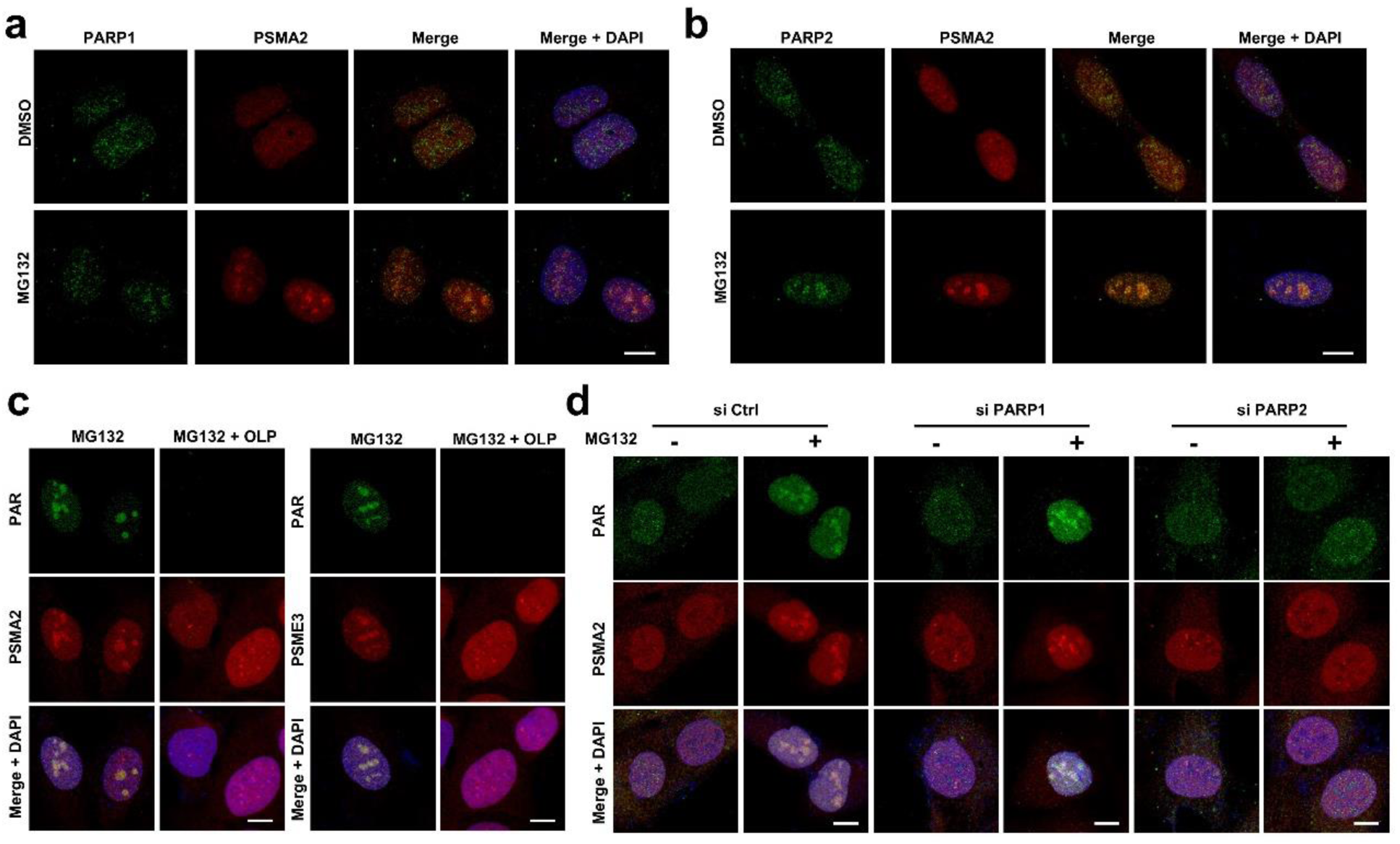
PAR-proteasome condensation depends on PARylation by PARP2. (a) and (b) Confocal microscopic imaging of PARP1 (a) and PARP2 (b) in MG132-treated U2OS cells. DMSO or MG132 (20 μM) was incubated with cells for 4 h. (c) Confocal microscopy of PAR-proteasome condensation in MG132-stimulated U2OS cells without and with pre-treatment by OLP. U2OS cells pre-treated without and with OLP (1 μM) for 0.5 h were incubated with MG132 (20 μM) for 4 h, followed by staining and imaging. (d) Confocal imaging of PAR-proteasome condensation in U2OS cells without and with PARP1 or PARP2 knockdown. U2OS cells without and with PARP1 or PARP2 knockdown by siRNAs were subjected to 4-h incubation in the absence and presence 20 μM MG132 and confocal microscopy. Scale bars, 10 µm.

Moreover, PAR-proteasome co-condensates were imaged in cells without and with exposure to PARP inhibitor OLP before MG132 stimulation. As shown by confocal microscopy, pre-incubation with OLP abolished the foci generation for both PAR and proteasome subunit PSMA2 and PSME3 after proteasomal inhibition (Fig. 3c), supporting necessity of PARylation catalytic activity for the condensation of PAR-proteasome. Then, U2OS cells with knockdown of PARP1 or PARP2 by siRNA were prepared for confocal microscopic studies of PAR-proteasome condensation (Fig. S3). The captured images indicated that unlike PARP1 knockdown with little effects on the development of PAR-proteasome co-condensates upon MG132 treatment, PARP2 knockdown blocks the generation of PAR or proteasome condensates in the presence of MG132 (Fig. 3d). Taken together, these results demonstrate that PARP2 is critical for the PAR-proteasome condensation formation.

### Ubiquitin chains are co-localized with PAR condensates

RAD23B, a shuttle factor for the ubiquitin-proteasome pathway, was next analyzed by confocal microscopy in U2OS cells as it was found to be necessary for proteasome condensation under hyperosmotic stress or nutrient starvation.^49,53^ Confocal images indicated co-localization of RAD23B with PAR condensates in MG132-stimulated cells (Fig. 4a). However, knockdown of RAD23B by siRNA in U2OS cells results in no changes to the development of PAR-proteasome co-condensates according to confocal imaging (Fig. 4b, S4a). These results revealed that different from previously observed phenomena regarding RAD23B-dependent proteasome condensation, PAR-proteasome condensation is independent of RAD23B.

**Figure 4.**
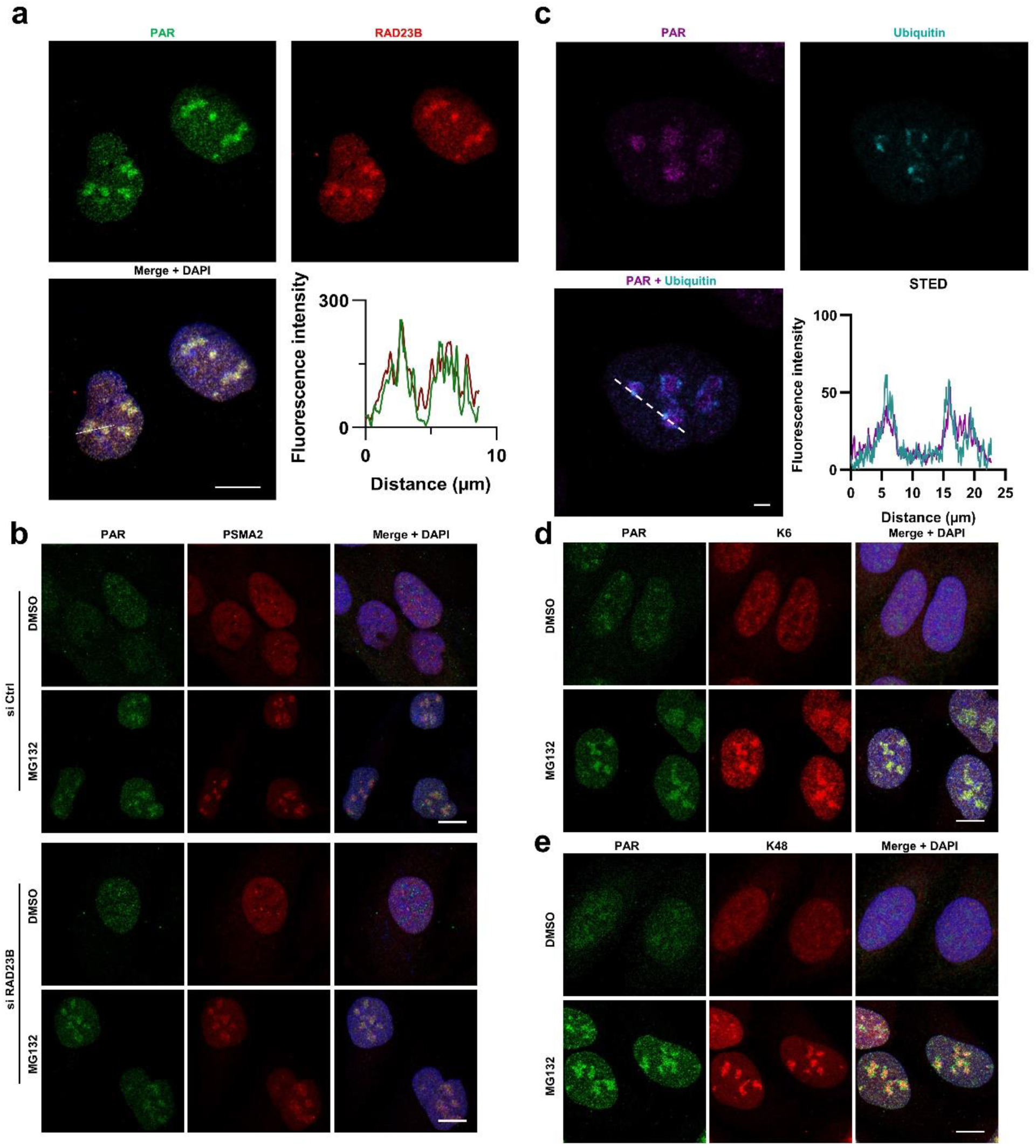
Co-localization of PAR condensates with ubiquitin chains. (a) Confocal microscopy of PAR and RAD23B in U2OS cells stimulated with MG132 (20 μM, 4 h). Plots at the bottom right show fluorescence signals of PAR and RAD23B from condensates indicated by the dashed line. Scale bars, 10 μm. (b) Confocal imaging of PAR-proteasome condensation in U2OS cells without and with RAD23B knockdown. Cells without and with RAD23B knockdown by siRNA were incubated for 4 h in the presence of DMSO or 20 μM MG132 prior to staining and imaging. Scale bars, 10 μm. (c) STED imaging of PAR and ubiquitin in MG132-treated U2OS cells. Following 4-h incubation with 20 μM MG132, cells were stained and analyzed by STED microscopy. Plots at the bottom right represent fluorescence intensities of PAR and ubiquitin from condensates indicated by the dashed line. Scale bars, 2.5 μm. (d) and (e) Confocal microscopic analysis of PAR and K6-linked ubiquitin chains (d) and PAR and K48-linked ubiquitin chains (e) in U2OS cells treated for 4 h with 20 μM MG132. Scale bars, 10 μm.

SUMO and ubiquitin were next examined for participation in PAR-proteasome condensation given their involvement in proteasome-regulated protein homeostasis.^54–57^ Immunoblots showed little changes in cellular levels of SUMO chains before and after incubation with proteasome inhibitor MG132, BTZ, and CFZ (Fig. S4b). Despite observation of liquid droplet-like structures for SUMO in U2OS cells upon MG132 treatment, no apparent co-localization of SUMO droplets with PAR condensates was seen in nucleus (Fig. S4c). By contrast, the levels of pan-ubiquitin, K6-, K48-, and K63-linked ubiquitin chains as measured by immunoblots were significantly increased after suppressing proteasomal activity by MG132, BTZ, and CFZ (Fig. S2b, S4d). And ubiquitin chains foci were observed in the nucleus of MG132-treated cells under confocal microscope, which exhibited a high degree of co-localization with PAR condensates (Fig. S4e). Moreover, the co-localization of PAR and ubiquitin foci was verified by super-resolution stimulated emission depletion (STED) microscopy, supporting direct interactions between PAR and ubiquitin chains within condensates (Fig. 4c). Also, confocal imaging analysis revealed that K6- and K48-linked ubiquitin chains formed condensates co-localized with PAR foci upon MG132 stimulation, whereas no condensation for K63-linked ubiquitin chains was detected (Fig. 4d, 4e, S4f). These data together are in support of co-localization of ubiquitin and PAR condensates.

### PAR-ubiquitin chain interaction and requirement of K6-linked ubiquitin chains for PAR condensation

We next corroborated binding interactions between K6- or K48-linked ubiquitin chains and PAR polymers. Recombinant human PARP2 or auto-PARylated PARP2 was mixed with K6- or K48-linked ubiquitin tetramer. Subsequent co-immunoprecipitation analysis using an anti-PARP2 monoclonal antibody demonstrated that tetra-K6 and tetra-K48 ubiquitin chains can directly bind to PARylated PARP2, but not to non-modified PARP2 (Fig. 5a, 5b). Furthermore, co-immunoprecipitation studies of purified PAR polymers mixed with the ubiquitin tetramers using an anti-PAR monoclonal antibody also supported direct binding between PAR polymers and ubiquitin chains (Fig. 5c, 5d). The PAR-ubiquitin chain interaction was also examined in the cellular context by PAR pulldown using U2OS cell lysates. Immunoblot analysis of the PAR pulldown samples indicated presence of ubiquitin chains including K6- and K48-linked ubiquitylation as well as proteasome subunits PSMA2 and PSMB5 in cells stimulated with MG132 (Fig. S5a). Additionally, sandwich ELISA assays revealed that auto-PARylated PARP2 binds tightly to K6- and K48-linked ubiquitin tetramers, whereas non-modified PARP2 displays little binding to the tetra ubiquitin chains under the same conditions (Fig. S5b). Along with above imaging results, these data support direct interactions between PAR and ubiquitin chains.

**Figure 5.**
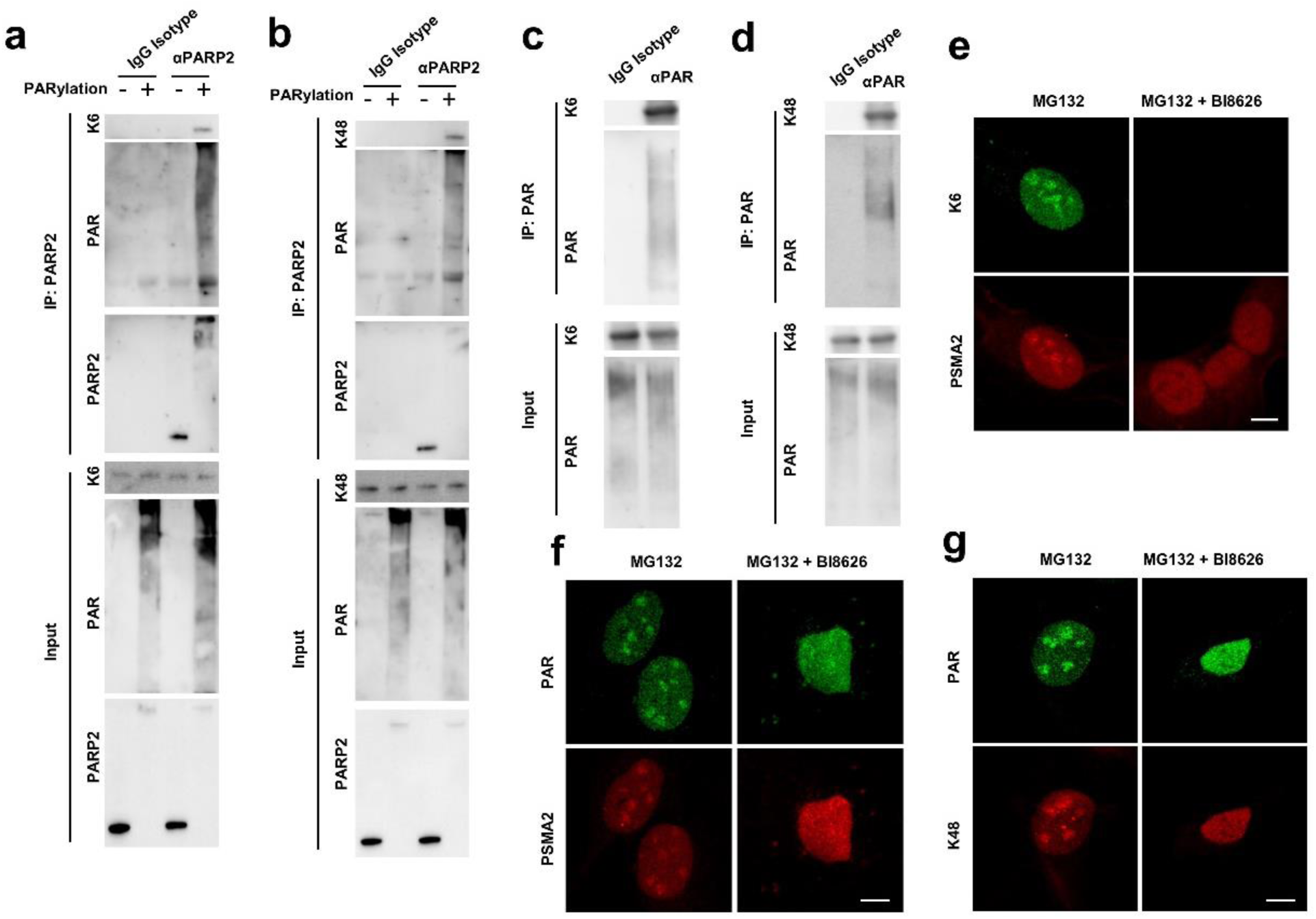
PAR binds to ubiquitin chains and requires K6-linked ubiquitin chains for condensation. (a) and (b) Co-immunoprecipitation of purified PARP2 and PARylated PARP2 mixed with tetra-K6 ubiquitin chains (a) or tetra-K48 ubiquitin chains (b). PARP2 was precipitated by an anti-PARP2 monoclonal antibody, followed by immunoblot analysis. (c) and (d) Co-immunoprecipitation of purified PAR incubated with tetra-K6 ubiquitin chains (c) or tetra-K48 ubiquitin chains (d). PAR was precipitated by an anti-PAR monoclonal antibody, followed by immunoblot analysis. (e) Confocal microscopy of K6-linked ubiquitin chains and PSMA2 in U2OS cells treated without and with BI8626 (10 μM) for 4 h in the presence of 20 μM MG132. (f) Confocal images of PAR and PSMA2 in MG132-stimulated U2OS cells without and with 10 μM BI8626 treatment. (g) Confocal microscopic analysis of PAR and K48-linked ubiquitin chains in MG132-treated U2OS cells without and with BI8626 (10 μM). Scale bars, 10 μm.

The impact of ubiquitin chains on PAR-proteasome condensation was then evaluated using a small-molecule inhibitor BI8626 specific for a major E3 ligase HUWE1 containing UBA and WWE domains.^58–61^ Immunoblots showed that relative to MG132-treated cells with accumulated K6- and K48-linked ubiquitylation, cells receiving both MG132 and BI8626 were characterized by substantially reduced K6-linked ubiquitin chains and mostly unaffected levels of K48-linked ubiquitylation (Fig. S5c). Importantly, confocal imaging of cells stimulated by MG132 revealed that additions of BI8626 inhibitor resulted in elimination of K6-link ubiquitylation signals and dissolution of condensates of PAR, proteasome subunits PSMA2 and PSME3, K48-linked ubiquitin chains, and PARP2 in nucleus (Fig. 5e, 5f, 5g, S5d, S5e). These results indicated that K6-linked ubiquitylation catalyzed by HUWE1 is indispensable for PAR-proteasome condensation upon proteasomal inhibition.

### Co-localization of PAR-proteasome condensates with stalled DNA replication forks

To unravel the cellular function of PAR-proteasome condensation, we delved into DNA replication in cells considering the association of proteasome activity with DNA replication as well as the detection of upregulated levels of p-H2AX, an early marker for DNA damage, upon exposing to proteasome inhibitors (Fig. S2b).^62,63^ Results from 5-ethynyl-2ʹ-deoxyuridine (EdU) incorporation assays revealed significant inhibition of DNA synthesis in U2OS cells following proteasome inhibition by MG132, BTZ, and CFZ (Fig. 6a). Moreover, DNA fiber assays using 5-iodo-2ʹ-deoxyuridine (IdU) illustrated that in comparison with the DMSO-treated control group, cells after incubation with proteasome inhibitors synthesized new DNA strands with overall reduced lengths (Fig. 6b). These data together indicated that DNA replication forks were stalled in cells receiving protease inhibitors.

**Figure 6.**
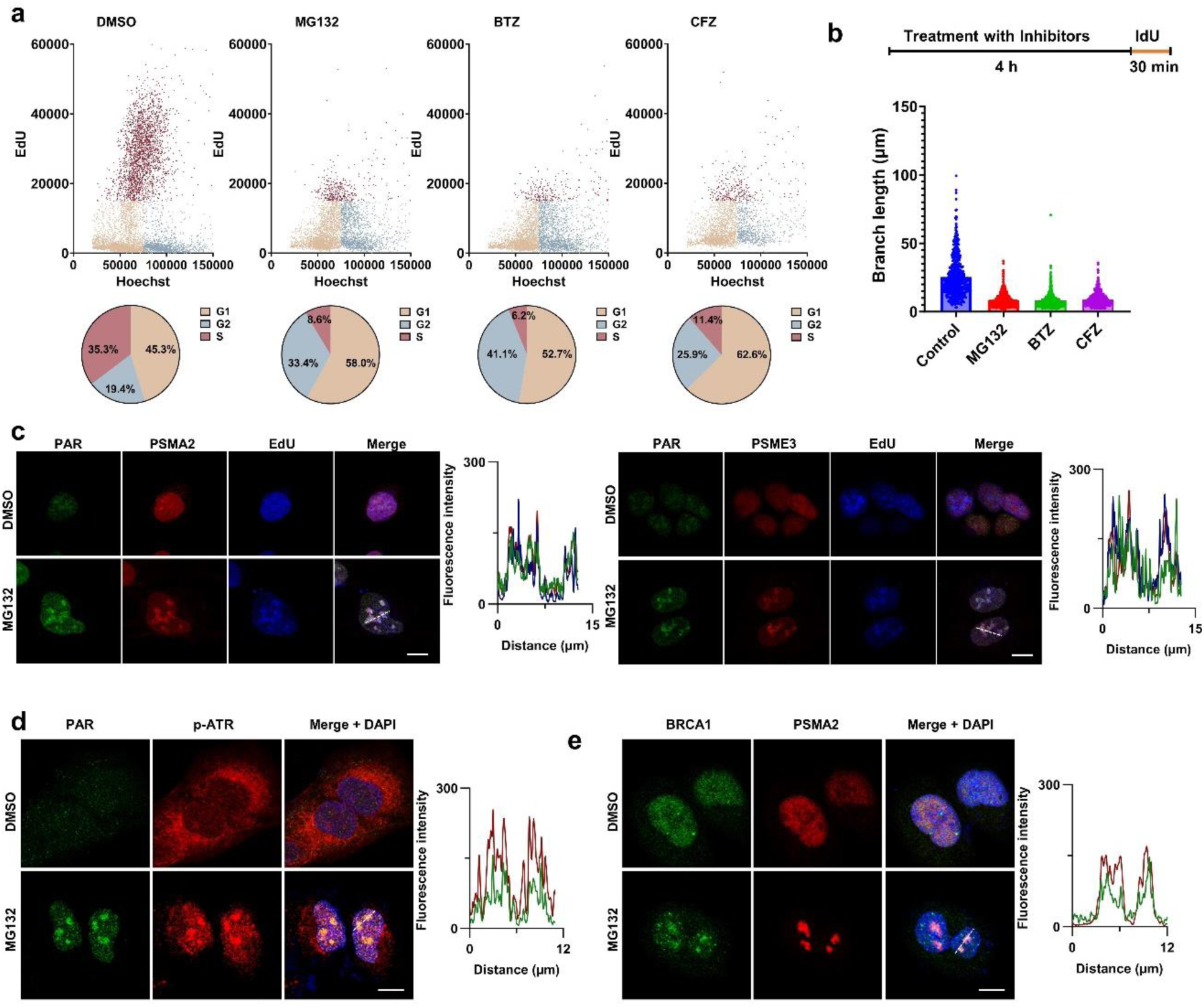
PAR-proteasome condensates are co-localized with stalled DNA replication forks. (a) EdU incorporation (top panel) and cell-cycle distribution (bottom panel) in U2OS cells after 4-h treatment with DMSO, MG132 (20 μM), BTZ (10 μM), or CFZ (10 μM). (b) Schematic of the DNA fiber assay and length of IdU-labeled tracks in U2OS cells without and with treatment by MG132, BTZ, and CFZ. (c) Confocal images of PAR, proteasome subunits, and incorporated EdU in U2OS cells treated with DMSO or MG132. Plots next to each image panel represent fluorescence signals of PAR, respective protease subunit, and EdU from condensates indicated by dashed lines. (d) and (e) Confocal microscopy of PAR and p-ATR (d) and BRCA1 and PSMA2 (e) in U2OS cells without and with MG132 treatment. Plots next to each image panel show fluorescence intensities of PAR and p-ATR (d) or BRCA1 and PSMA2 (e) from condensates indicated by dashed lines. Scale bars, 10 μm.

We then examined subcellular locations of DNA replication forks by confocal microscopy. Incorporated EdU-derived foci were observed under confocal microscope in the nucleus of cells treated by MG132, BTZ, and CFZ (Fig. S6a). Strikingly, confocal imaging analysis indicated that EdU foci found in MG132-stimulated cells were co-localized with PAR-proteasome condensates (Fig. 6c). Furthermore, co-localization was observed in cells with MG132 for PAR-proteasome condensates and markers of stalled DNA replication forks, including phosphorylated ATR (p-ATR) and BRCA1 (Fig. 6d, 6e). In addition, confocal images revealed multiple p-H2AX foci in nucleus that were strongly co-localized with PAR foci among cells incubated with MG132, BTZ, and CFZ (Fig. S6b). These results collectively support that PAR-proteasome condensates are co-localized with stalled replication forks upon proteasomal inhibition, implicating involvement of PAR-proteasome condensation in replication fork stalling-induced DNA damage response.

### PAR-proteasome condensation sustains genomic stability through protection of stalled replication forks

Next, the function of PAR-proteasome condensation was probed using HUWE1 inhibitor BI8626 and PARP inhibitor OLP that block the generation of PAR-proteasome condensates in MG132-treated cells (Fig. 3c, 5f, S5d). EdU incorporation analysis showed that in comparison to MG132-stimulated U2OS cells with markedly lower levels of DNA synthesis than the DMSO-treated group, cells receiving MG132 along with BI8626 were partially relieved from inhibition of DNA synthesis (Fig. 7a), indicating attenuation of DNA replication by PAR-proteasome condensation. Immunoblots of cells incubated in the absence and presence of proteasome inhibitors (MG132, BTZ, and CFZ), OLP, and/or BI8626 revealed that relative to groups with proteasome inhibitors only, treatments combining the proteasome inhibitor with OLP and/or BI8626 led to increased levels of the p-H2AX DNA damage marker (Fig. 7b). Moreover, terminal deoxynucleotidyl transferase dUTP nick-end labeling (TUNEL) assay analysis illustrated significantly enhanced levels of DNA lesion for cells treated by MG132 in combination with OLP and/or Bl8626 over ones by MG132 alone, consistent with the immunoblot data (Fig. 7b, 7c). These results indicated that upon proteasomal inhibition, the formed PAR-proteasome condensates facilitate stalling and protection of DNA replication forks.

**Figure 7.**
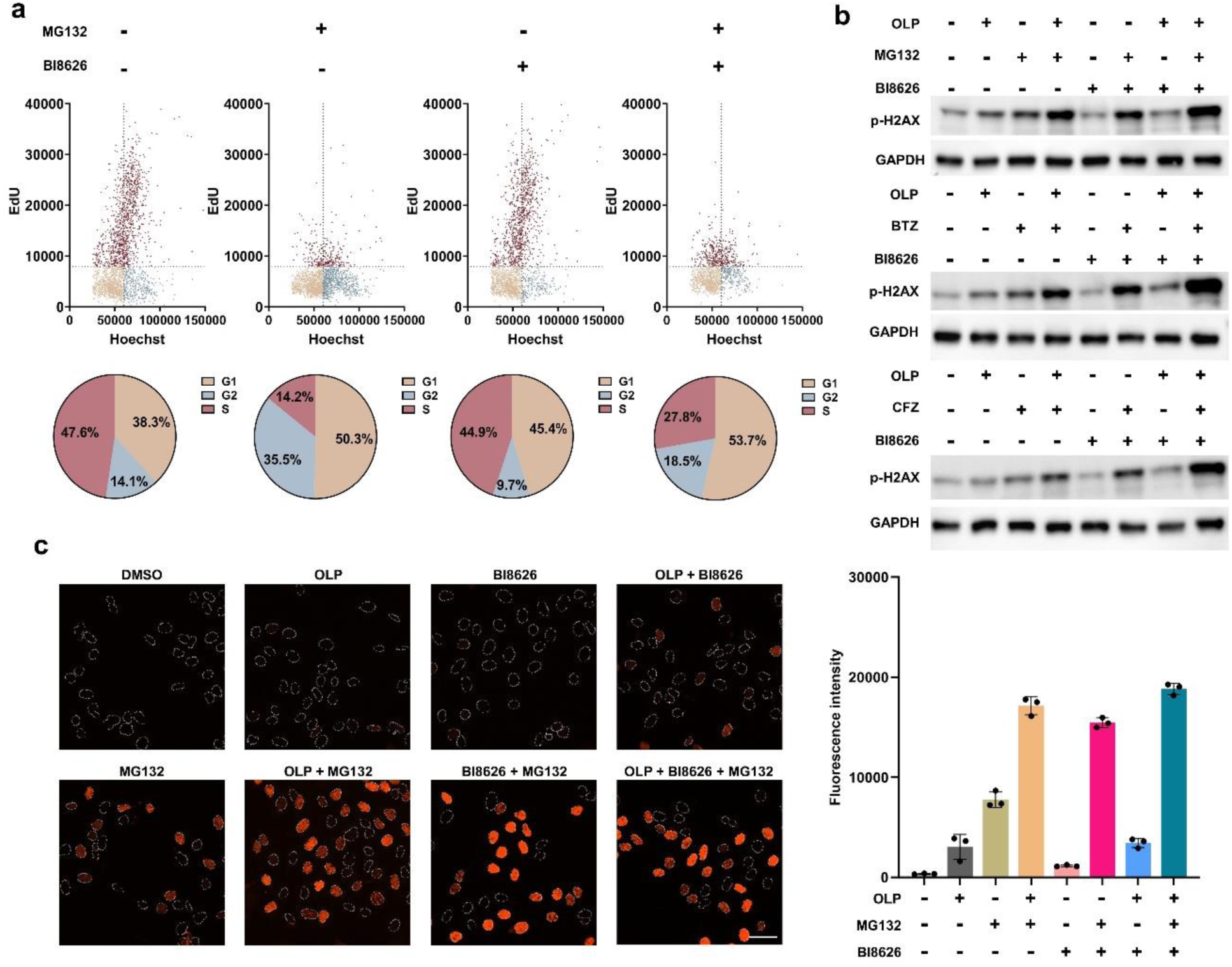
Blocking PAR-proteasome condensation leads to increased DNA damage. (a) EdU incorporation (top panel) and cell-cycle distribution (bottom panel) in U2OS cells after treating for 4 h in the absence and presence of 20 μM MG132 and/or 10 μM BI8626. (b) Immunoblot analysis of p-H2AX of U2OS cells incubated without and with proteasome inhibitors (MG132: 20 μM; BTZ and CFZ: 10 μM), OLP (1 μM), and/or BI8626 (10 μM). (c) Fluorescence imaging of DNA damages in the nucleus of U2OS cell by TUNEL assays following treatment in the absence and presence of proteasome inhibitors, OLP, and/or BI8626. Right panel: fluorescence intensity for cells under each condition. Scale bars, 50 μM.

## Discussion

This study uncovers a unique form of nuclear condensation triggered by proteasomal inhibition. It features macromolecular assemblies of PAR, proteasome, and ubiquitin. Condensing PAR polymers arise from PARP2 catalysis and directly interact with ubiquitin chains within co-condensates. K6-linked ubiquitin chains are essential for PAR-proteasome condensation. In response to proteasomal stress, PAR-proteasome-ubiquitin chain co-condensation functions by retarding DNA replication and confining stalled replication forks in condensates to maintain genome integrity (Fig. 8). Given its occurrence in multiple types of cells following inhibition of proteasome activity, PAR-proteasome-ubiquitin chain condensation may represent a general protective mechanism in the nucleus of mammalian cells under stressed conditions.

**Figure 8.**
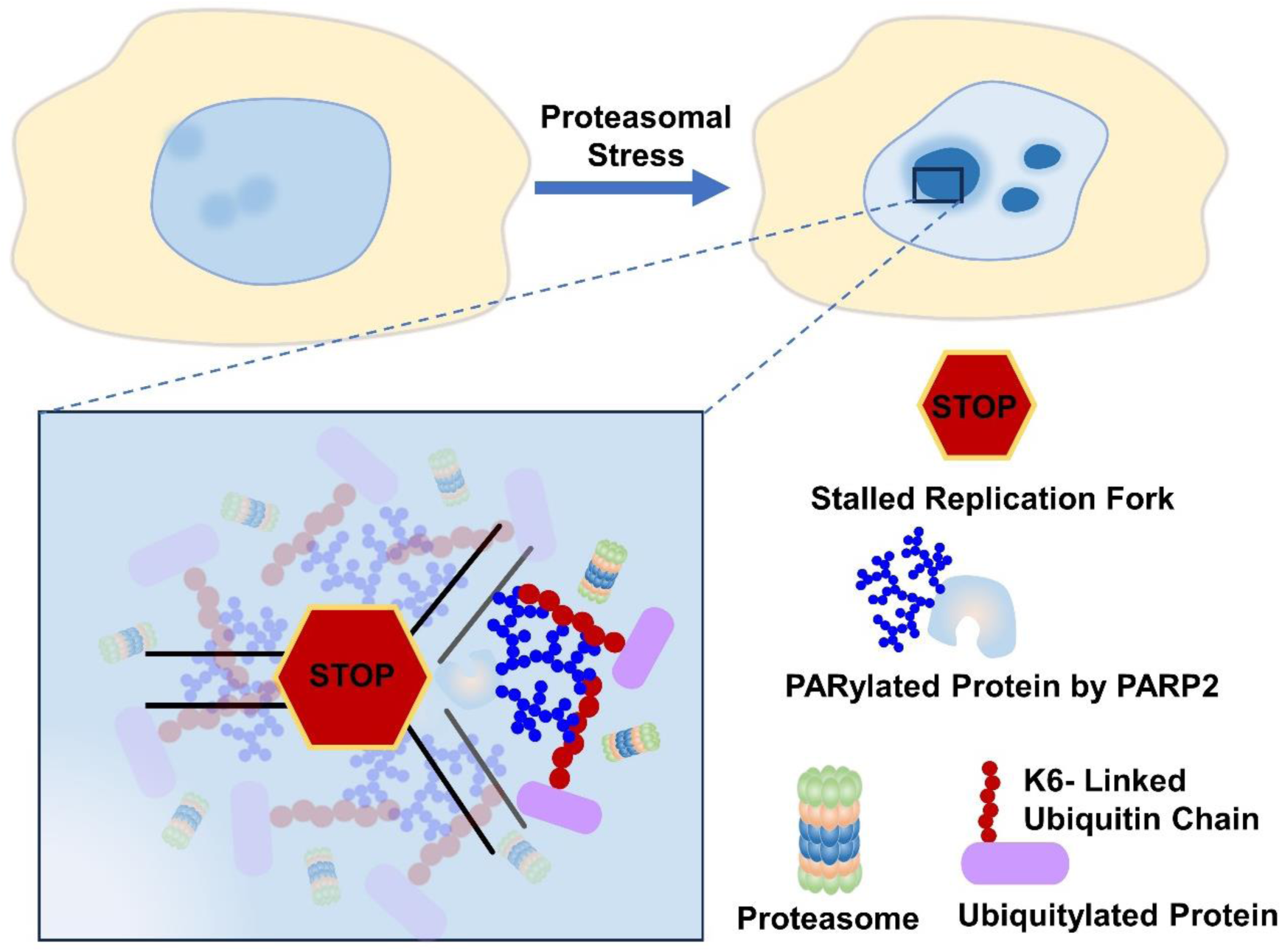
Schematic representation of the cellular function of PAR-proteasome-ubiquitin chain condensates formed during proteasomal stress.

PAR synthesized by PARP1 or PARP5a was recently found to participate in condensation of nucleosomes and proteins with intrinsically disordered regions as well as stress granule formation.^45–48^ Recent reports also showed RAD23B-dependent condensation of proteasome in response to osmotic shock or amino acid starvation.^49,53^ Completely different from previous findings, this work revealed that PARP2-catalyzed PARylation undergoes co-condensation with proteasome and ubiquitin chains upon inhibiting proteasome activity. And RAD23B is nonessential for PAR-proteasome condensation despite co-localization of RAD23B with PAR condensates. Therefore, our findings present a new class of biomolecular condensates with pivotal functions. The observation of PAR and proteasome in condensates implies direct interactions between PAR and proteasome. In fact, previous proteomic studies identified proteasome subunits as putative PAR binders.^20,64^

While both PARP1 and PARP2 catalyze PARylation in nucleus, only PARP2 was co-localized with PAR-proteasome condensates and required for the condensation formation following inhibition of proteasome activity. These findings suggest differential capabilities of PAR synthesized by PARP1 and PARP2 in promoting condensation with other macromolecules. In contrast to PARylation by PARP1, PARP2-derived PAR polymers usually carry more branches,^15^ which may improve binding affinity and/or specificity of this multivalent scaffold toward PAR-binding proteins such as ubiquitin chains and proteasome subunits and consequently foster biomolecular condensation.

As conserved post-translational modifications, PARylation and ubiquitylation are recognized by a variety of proteins. Here our data for the first time demonstrate direct binding between PAR polymers and ubiquitin chains. Because PAR carries negative charges and ubiquitin chains are positively charged at physiological pH, these two polymeric structures are likely to form strong ionic interactions. The resulting PAR-ubiquitin chain complexes could serve as reversible scaffolds central to macromolecular assemblies. Upon inhibiting proteasome activity, upregulated PAR and ubiquitin chains may orchestrate recruitments of proteasome and other biomolecules and subsequent co-condensation to stabilize arising stalled replication forks, generating a potentially universal response to this stress.

In conclusion, our study uncovered nuclear co-condensates of PAR with proteasome and ubiquitin chains in cells experiencing proteasomal inhibition as well as binding of PAR to ubiquitin chains. PARP2 and K6-linked ubiquitin chain are required for the co-condensation. By protecting stalled DNA replication forks resulted from proteasomal stress, PAR-proteasome-ubiquitin chain co-condensates play a key role in sustaining genome stability, shedding new light on biomolecular condensation in stressed cells.

## Methods

### Cell culturing

U2OS, A2780, and HEK293T cells were cultured in DMEM medium supplemented with 10% fetal bovine serum (FBS) and 1% penicillin/streptomycin. LNCaP and J774A.1 cells were grown in RPMI-1640 medium supplemented with 10% FBS. All cells were maintained in 37°C incubators with 5% CO_2_.

### Compound screening

A group of 60 compounds were screened for inducing liquid-liquid phase separation (LLPS) of PAR in U2OS cells using an anti-PAR monoclonal antibody (clone: 10H) via immunofluorescence imaging. Briefly, U2OS cells were seeded in 96-well plates (5000 cells per well). After overnight incubation and attachment to the bottom of plates, cells were treated with various types of compounds for up to 4 h. Fixation and immunostaining were performed as described in the below subsection. LLPS was quantified as the ratio of the phase separation area to the nuclear area in individual cells in µm^2^. Z-score was calculated for each measurement with the formula, Z-score=(value-mean)/standard deviation. Each compound treatment was assessed across 3-6 independent experiments with 500-1000 cells analyzed per condition. Images were collected using Cytation5 and analyzed using Image J.

### Immunofluorescence staining and confocal microscopy

Cells were fixed with 4% paraformaldehyde in PBS for 15 min, followed by rinsing and permeabilization with 0.1% Triton X-100 in PBS for 10 min. After incubating with 10% FBS in PBS at room temperature for 1 h, cells were stained with primary antibodies in 10% FBS in PBS overnight at 4°C. After washing with PBS three times, cells were then incubated with secondary antibodies at room temperature for 1 h. Nuclear staining was performed with DAPI or Hoechst. Cells were next rinsed with PBS and imaged using a ZEISS LSM 880 Airyscan microscope equipped with a 63× objective lens.

### STED microscopy

Imaging was performed using a Leica Stellaris 8 microscope (Leica Microsystems) equipped with a Leica scanhead, a white-light laser for excitation, 592-nm and 775-nm depletion lasers, Leica HyD S and HyD X counting detectors, and an HC PL APO 86×/1.20 motCORR STED water immersion objective. STED microscope alignment and performance were verified before each experiment using fluorescent beads. Briefly, 20-nm fluorescent microspheres (F8782, Thermo Fisher Scientific) were immobilized on poly-D-lysine-coated (A3890401, Thermo Fisher Scientific) #1.5 glass coverslips (Zeiss) mounted on microscope slides. After automatic alignment of the Stellaris 8 in LAS X 4.8.2 (Leica Microsystems), manual alignment was performed to ensure perfect spatial overlap of the excitation and depletion beams using the bead samples under the same imaging conditions (Fig. S7).

For Alexa 488, excitation was performed at 491 nm, and emission was collected between 505 nm and 565 nm. A constant-wavelength 592-nm laser was used for depletion. For Alexa 647, excitation was performed at 638 nm, and emission was collected between 652 nm and 750 nm. A pulsed 775-nm laser was used for depletion. Temporal gating was applied from 1 to 8 ns. Laser powers were kept constant to ensure comparability across images. The white-light laser was set to 80% total power with 4% for Alexa 488 excitation and 7% for Alexa 647 excitation. The 592-nm laser was set to 100% power, using 75% for depletion. The 775-nm laser was also at 100% power, with 50% for depletion. The pixel size was 33 nm (1024 × 1024 pixels), and the pixel dwell time was 1.575 µs. Four-line accumulations were collected for all images.

### RNA interference and transfection

siRNA oligos for human PARP1 (siRNA ID:10038), PARP2 (siRNA ID:111561), RAD23B (siRNA ID:12169), and non-targeting siRNA control (catalog number: 4390843) were purchased from Thermo Fisher Scientific. U2OS cells were transfected with siRNAs using Lipofectamine RNAiMAX (Thermo Fisher Scientific) and then collected for analysis 36 h after transfection.

### Immunoblot analysis

Cells were lysed in RIPA buffer containing protease and phosphatase inhibitor cocktail on ice for 30 min with rotation. Following centrifugation at 12000×g for 10 min at 4°C, supernatants were collected. Protein concentrations were determined using BCA protein assay kits (Thermo Fisher Scientific). Cell lysates were electrophoresed in 4-20% Express Plus PAGE gels (GenScript) and transferred to polyvinylidene difluoride membranes (Bio-Rad). After blocking with 5% BSA and incubating with antibodies and SuperSignal West Pico PLUS chemiluminescent substrate (Thermo Fisher Scientific), membranes were imaged using a ChemiDoc touch imaging system (Bio-Rad).

### Human PARP2 expression and purification

The expression and purification of human His_6_-tagged full-length PARP2 were performed as previously described with minor modifications.^5^ Briefly, BL21 (DE3) cells were transformed with the constructed pET-28a vector expressing full-length human PARP2 and grown in LB broth with kanamycin (50 µg mL^−1^) at 37°C in shaker flasks (250 rpm). After 600-nm optical density reached 0.6-0.8 and subsequent additions of 0.1 mM ZnSO_4_ and 0.5 mM isopropyl β-D-thiogalactoside, cells were induced for PARP2 expression at 16°C for 24 h. Cell pellets were collected by centrifugation at 4550×g for 30 min at 4°C, resuspended in lysis buffer (25 mM HEPES pH 8.0, 500 mM NaCl, and 1 mM phenylmethylsulfonyl fluoride), and lysed by a French Press (GlenMills, NJ). Cell lysates were centrifuged at 15000×g for 60 min at 4°C. Supernatants were filtered with 0.45 µm membranes, followed by loading onto a gravity-flow column with nickel-nitrilotriacetic acid agarose beads (Thermo Fisher Scientific). The column was washed with 25 mM HEPES pH 8.0 with 500 mM NaCl and 20 mM imidazole and eluted with 25 mM HEPES pH 8.0 with 500 mM NaCl and 400 mM imidazole. After adding equal volume of 50 mM Tris pH 7.0 with 1 mM EDTA and 0.1 mM DTT, eluate was injected into a HiTrap heparin column. Following column washing with 50 mM Tris pH 7.0 with 1 mM EDTA, 0.1 mM DTT, and 250 mM NaCl, PARP2 was eluted with 50mM Tris pH 7.0 with 1 mM EDTA, 0.1 mM DTT, and 1 M NaCl and concentrated using 30-kDa cutoff centrifugal filters. The PARP2 concentration was determined by a NanoDrop 2000C UV-Vis spectrophotometer (Thermo Fisher Scientific).

### Auto-PARylation of PARP2

Auto-modification of PARP2 was performed as previous report.^5^ Briefly, 3 μM PARP2 was pre-incubated in 100 mM Tris-HCl at pH 8.0 with 50 mM NaCl, 10 mM MgCl_2_, 100 ng μL^−1^ activated DNA, and 1 mM DTT at 30°C for 30 min, followed by 4-h incubation at 30°C in the absence and presence of 300 μM NAD^+^. After stopping reactions with 100 μM olaparib, PARP2 and PARylated PARP2 were analyzed by immunoblots using an anti-PAR monoclonal antibody (clone: 10H).

### Co-immunoprecipitation

U2OS cells were lysed in RIPA buffer containing protease and phosphatase inhibitor cocktail on ice for 30 min with rotation. Supernatants were collected after centrifugation at 16000×g for 10 min. Protein concentrations were determined using BCA protein assay kits. For co-immunoprecipitation of PAR-associated proteins, an anti-PAR monoclonal antibody (clone: 10H) and protein G agarose beads (GenScript) were added into the cell lysates and incubated at 4°C for 4 h. After washing beads with RIPA buffer five times, proteins were eluted at 80°C for 10 min in LDS buffer (GenScript).

For *in vitro* co-immunoprecipitation of PARP2-binding proteins, 50 nM purified PARP2 or PARylated PARP2 was mixed with 100 nM tetra-K6 or tetra-K48 ubiquitin chains. For *in vitro* co-immunoprecipitation of PAR-interacting proteins, 200 nM purified PAR (Bio-Techne) was mixed with 200 nM tetra-K6 or tetra-K48 ubiquitin chains. The mixtures were incubated at 4°C for 1 h, followed by additions of protein G agarose beads together with an anti-PARP2 monoclonal antibody (clone: F-3) or anti-PAR monoclonal antibody (clone: 10H) and 4-h incubation at 4°C. After washing beads with RIPA buffer three times, samples were analyzed via immunoblots.

### Label-free quantitative mass spectrometry

U2OS cells were treated with DMSO or MG132 (20 µM) for 4 h and lysed in RIPA buffer containing protease and phosphatase inhibitor cocktail on ice for 30 min with rotation. The supernatants were collected after centrifugation at 12000×g for 15 min. Anti-PAR immunoprecipitation was performed at 4°C overnight using protein G agarose beads with an anti-PAR monoclonal antibody (clone: 10H) or an IgG isotype control. After washing the beads with RIPA buffer four times, proteins were eluted with SDS-PAGE loading buffer and reduced with 10 mM tris(2-carboxyethyl)phosphine for 0.5 h at 55°C. The samples were then alkylated with 18.75 mM iodoacetamide for 20 min in the dark at room temperature, followed by additions of six volume of pre-chilled acetone, vortexing, and overnight precipitation at −20°C. After centrifugation at 8000xg for 10 min at 4°C, pellets were dried at room temperature for 5 min and resuspended in 12.5 μL denaturing buffer (100 mM Tris pH 8.5 and 8 M urea) at room temperature for 15 min on a shaker at 800 rpm. Samples were subsequently diluted to bring the concentration of urea to 1 M with 100 mM Tris pH 8.5 and digested with 0.1 µg trypsin (Promega V5280) at 37°C for 18 h on a shaker at 225 rpm. Samples were acidified to 0.66% TFA and centrifuged at 16000xg for 5 min at 4°C. Supernatants were collected and extracted with 2 µg-capacity C18 ZipTips (Millipore ZTC18M096). The dried desalted peptides were resuspended in 8 µL of 0.1% TFA and 2 µL samples were injected onto an UltiMate 3000 UHP liquid chromatography system (Thermo Fisher Scientific).

Peptides were trapped on an Acclaim PepMap 100 C18 trapping column (5 μm, 0.1×150 mm; Thermo Fisher Scientific) using a loading pump operating at 10 µL min^-1^ with 2% acetonitrile and 0.1% TFA in water. After desalting for 3 min, peptides were separated on a 200 cm μPAC column (Thermo Fisher Scientific) using a linear gradient condition (1-4 min, 5-8% solvent B; 4-70 min, 4-30% solvent B; 70-90 min, 30-55% solvent B; 90-112 min, 55-97% solvent B; 112-120 min, 97% solvent B) at a flow rate of 700 nL min^-1^. Gradient solvent A, 0.1% formic acid; solvent B, 80% acetonitrile and 0.1% formic acid.

Eluted peptides were electrosprayed into an Orbitrap Fusion Lumos Tribrid mass spectrometer (Thermo Fisher Scientific) equipped with a Nanospray Flex ion source (Thermo Fisher Scientific) coupled to a FAIMS Pro Duo (Thermo Fisher Scientific). The instrument method performed by the Xcalibur v 4.7 software consisted of the following standard mass spectrometer parameters: spray voltage, 2.5 kV; heated capillary temperature, 300°C; S-Lens RF level, 30%; FAIMS total carrier gas flow, 4.0 L s^-1^; and FAIMS cooling gas flow, 5.0 L s^-1^. The instrument was operated in data-dependent acquisition mode. Full-scan mass spectra (375–1500 m/z, 60000 resolution) were detected in the orbitrap analyzer with ‘standard’ automatic gain control. FAIMS compensation voltages were set to −45 and −60 V. The monoisotopic precursor selection (MIPS) filter was set to peptides and candidate precursors above the 5.0e3 signal threshold were added to the dynamic exclusion list for 60 s with a 10-ppm mass window following a single selection for MS2. Precursor ions with charge states < +2 and > +6 were excluded. For every full scan, MS/MS scans were collected in the ion trap during a 1.5 s cycle time. Ions were isolated (1.6 m/z isolation width) via quadrupole for higher-energy collisional dissociation (HCD) fragmentation at 35% normalized collision energy. Fragment ion mass spectra were taken with the ion trap operating in ‘turbo’ mode with ‘automatic’ maximum injection time.

Thermo.Raw files were imported into Proteome Discoverer 3.0 (Thermo Fisher Scientific) using default parameters. The search engine Sequest HT was used for peptide assignment. Parameters for protein search and identification were defined as follows: database, Uniprot human proteome database; precursor mass tolerance, 10 ppm; fragment tolerance, 0.2 Da; digestion, trypsin with up to two missed cleavages; fixed modification, carbamidomethyl (C); variable modifications, oxidation (M) and ubiquitination remnant GG (K). Proteins were identified with at least two high confidence peptides with a 1% false discovery rate (FDR). Label-free precursor quantification was performed using Minora feature detector with protein abundances calculated based on summed abundances of peptides and *P* values derived from a t-test. To find PAR-interacting or PARylated proteins that were enriched in the MG132-treated condition, proteins were first identified that were significantly enriched over the IgG isotype control (at least 3-fold enriched with *P* value < 0.05). Then, abundance ratios of MG132-treated *vs* DMSO-treated samples were calculated for these putative PAR-interacting or PARylated proteins.

### ELISA analysis of binding of ubiquitin chains to PARylated PARP2

Sandwich ELISA was conducted with tetra-K6 or tetra-K48 ubiquitin chains as the capture reagent and an anti-PARP2 monoclonal antibody as the detection reagent. High-binding 96-well plates were coated with tetra-K6 or tetra-K48 ubiquitin chains (1 µg mL^-1^) overnight at room temperature. After washing with PBS containing 0.05% tween-20 (PBST), wells were blocked with 5% BSA in PBS for 4 h at room temperature. PARP2 or PARylated PARP2 at varied concentrations was added and incubated for 2 h at room temperature. Following three washes with PBST, goat anti-mouse IgG horseradish peroxidase (HRP) was added and incubated for 1 h at room temperature. Plates were then washed with PBST five times, followed by additions of QuantaBlu fluorogenic peroxidase substrates (Thermo Fisher Scientific). Fluorescence intensities (excitation, 325 nm; emission, 420 nm) were measured using a Synergy H1 hybrid multi-mode microplate reader.

### Cell cycle analysis

U2OS cells were plated in confocal dishes and incubated with DMSO, MG132 (20 µM), BTZ (10 µM), or CFZ (10 µM) for 3.5 h. After adding 10 µM 5-ethynyl-2′-deoxyuridine (EdU) and incubating for 30 min, cells were fixed with 4% paraformaldehyde in PBS at room temperature for 15 min. Cells were then washed with PBS three times, permeabilized with 0.25% Triton X-100 in PBS, and blocked with 5% BSA in PBS. Copper(I)-catalyzed azide-alkyne cycloaddition was performing for 30 min at 37°C in 100 mM Tris pH 8.5 with 5 mM CuSO_4_ and 10 µM AFDye647 Azide, and 10 mM L-ascorbic acid. Nuclei were identified via Hoechst 33342 staining. Images were collected using the Cytation5 cell imaging multi-mode reader at 20x magnification. Nuclei were designated as individual regions of interest (ROIs) using Fiji software (Image J). According to EdU and Hoechst signaling intensities, cells were classified into different phases. Approximately 5000 cells were analyzed for each treatment group.

### DNA fiber assay

U2OS cells were treated with DMSO, MG132 (20 µM), BTZ (10 µM), or CFZ (10 µM) for 4 h and then labelled with 250 µM 5-iodo-2′-deoxyuridine (IdU) for 30 min. Cells were subsequently harvested and placed on slides for attachment. After cell lysis and spreading, samples were fixed using a mixture of methanol and acetic acid (3:1 v/v) for 10 min. Following two washes with PBS and denaturing with 2.5 M hydrochloric acid for 1 h, slides were blocked with 5% BSA and 0.1% Triton X-100 in PBS. After staining with a mouse anti-5-bromo-2′-deoxyuridine (BrdU) monoclonal antibody and subsequently an Alexa Fluor 488 goat anti-mouse IgG antibody, samples were fixed with 4% paraformaldehyde in PBS for 10 min and rinsed with PBS. Images of 300 DNA fibers for each treatment group were taken using a Leica SP8 confocal microscope for analysis.

### TUNEL assay

The terminal deoxynucleotidyl transferase dUTP nick-end labeling (TUNEL) assay was performed using Click-iT Plus TUNEL assay kits (C10618, Thermo Fisher Scientific) by following the manufacturer’s protocol with minor modifications. Briefly, U2OS cells were seeded in 8-well dishes and treated without and with MG132 (20 µM), olaparib (1 µM), and/or BI8626 (10 µM) for 4 h. After fixation with 4% paraformaldehyde in PBS for 15 min and permeabilization with 0.25% Triton X-100 in PBS for 15 min at room temperature, cells were incubated with 100 µL TdT reaction mixtures (2 µL EdUTP, 4 µL TdT enzyme, and 94 µL TdT reaction buffer) at 37°C for 10 min. Following two washes with 3% BSA in PBS for 5 min each, cells underwent incubation in 100 µL click reaction buffer (90 µL Click-iT Plus TUNEL Supermix and 10 µL 10×Click-iT Plus TUNEL reaction buffer additive) at 37°C for 30 min. Counter-staining was performed using Hoechst 33342. Cells were imaged and signaling intensity was quantified using Fiji software by measuring integrated density within individual nuclei. Approximately 2500-4000 cells were analyzed for each treatment group.

### Statistical analysis

Data is shown as mean ± standard deviation from three or more biologically independent experiments. The significance of difference in groups was compared via a two-tailed unpaired Student’s t-test. Multiple comparisons were performed using adjusted *P* values. All statistical analyses were performed using GraphPad Prism.

## Supporting information

The supporting information includes supplementary methods and results.

## Data availability

All data supporting the findings of this study are included in the main text and supplementary figures.

## Author contribution

L.Z. and Y.Z. designed research. L.Z., Z.Z., T.R.O., A.S. and G.K. performed research. L.Z., Z.Z., T.R.O., P.G. and Y.Z. analyzed data. L.Z. and Y.Z. wrote the manuscript.

## Acknowledgments

The STED imaging was performed at USC Translational Imaging Center. This work was supported by National Institute of General Medical Sciences (NIGMS) of the National Institutes of Health (NIH) grant R35GM137901 (to Y. Z.).

