## Supporting Information for "PAR-Driven Condensation Maintains Stalled Replication Fork Stability"

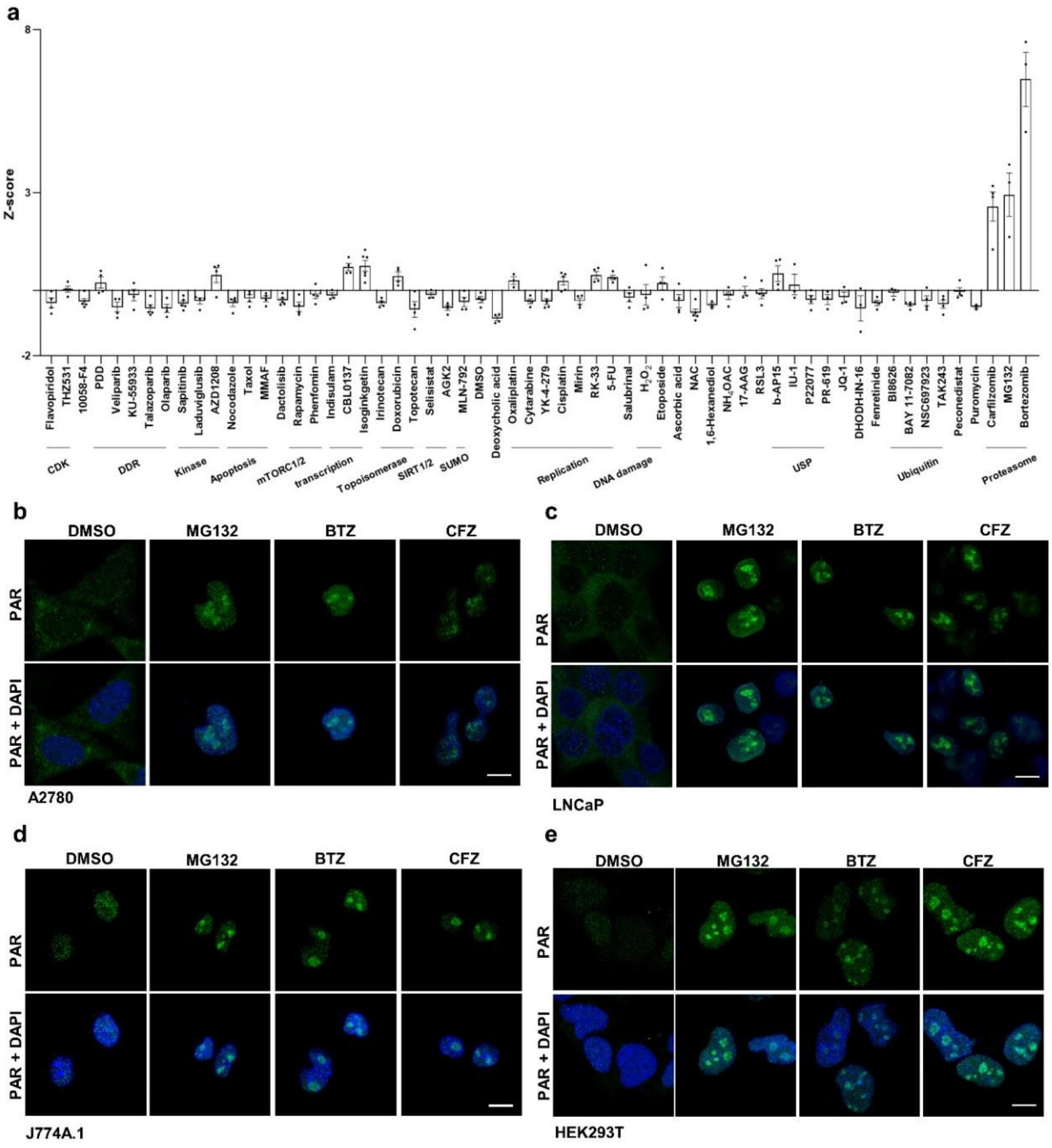


Figure S1. Proteasome inhibitors promote PAR condensation. (a) Z-scores of an in-house small-molecule library in driving liquid-liquid phase separation probed by an anti-PAR monoclonal antibody (clone: 10H). U2OS cells were incubated with compounds for 4 h and then stained for fluorescence imaging. Phase separation for 500-1000 cells under each condition was analyzed per replicate. CDK, cyclin-dependent kinases. DDR: DNA damage response. USP, ubiquitin-specific protease. (b)-(e) Immunofluorescence imaging of PAR condensation in A2780 (b), LNCaP (c), J774A.1 (d), and HEK293T (e) cells after 4-h treatment with MG132 (20 μM), BTZ (10 μM), or CFZ (10 μM). Scale bars, 10 μm.

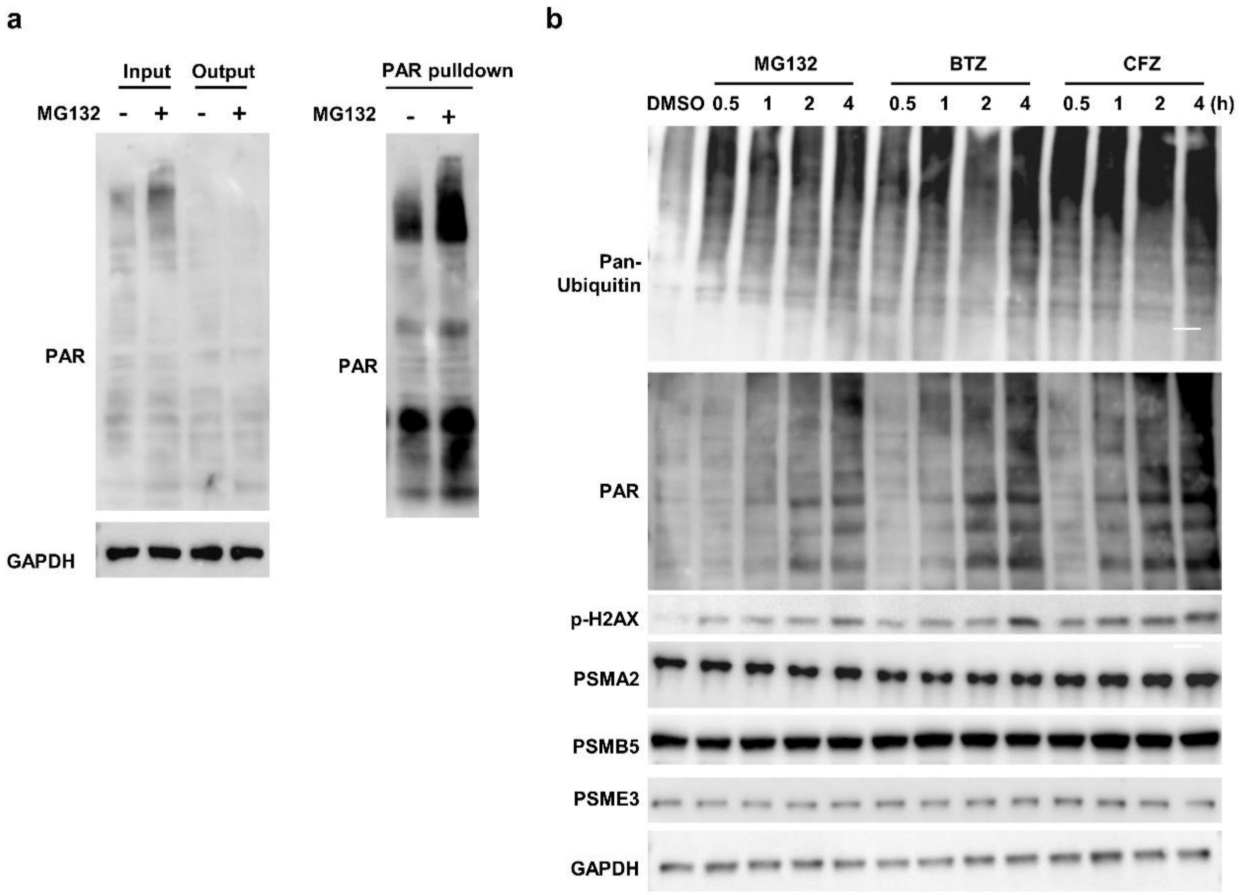


Figure S2. Immunoblot analysis of U2OS cells without and with protease inhibitor treatment. (a) Immunoblots of PAR pulldown samples from lysates of U2OS cells without and with 4-h treatment of 20 μM MG132. PAR was precipitated by an anti-PAR monoclonal antibody, followed by immunoblot analysis. (b) Immunoblots of U2OS cell lysates after up to 4-h treatment with DMSO, 20 μM MG132, 10 μM BTZ, or 10 μM CFZ.

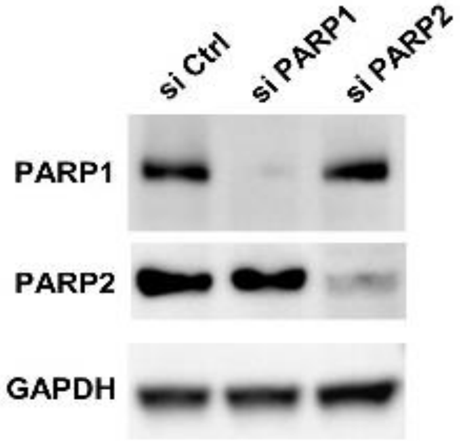


Figure S3. Immunoblots of PARP1 or PARP2 knockdown by siRNA in U2OS cells.

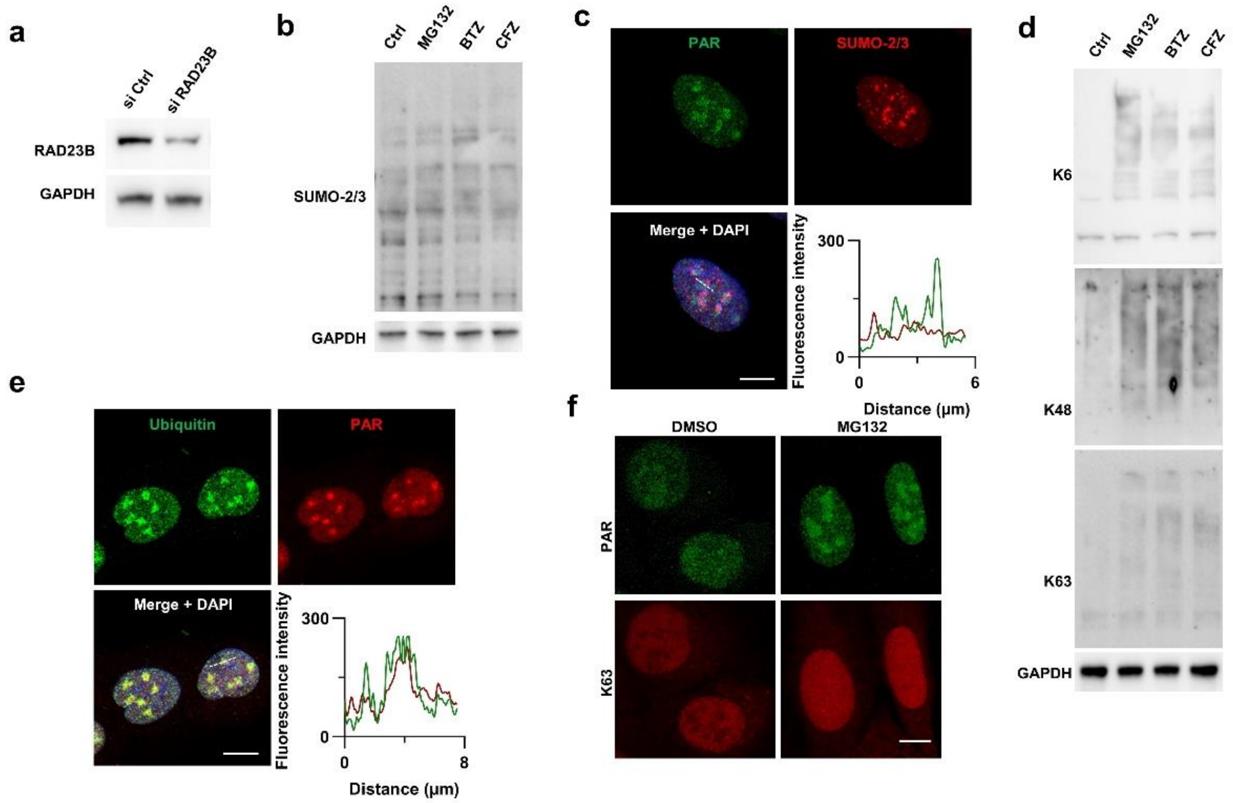


Figure S4. Accumulated ubiquitin chains are co-localized with PAR condensates. (a) Immunoblots of RAD23B knockdown by siRNA in U2OS cells. (b) Immunoblots of SUMO-2/3 in U2OS cells after 4-h treatment without and with MG132 (20 μM), BTZ (10 μM), or CFZ (10 μM). (c) Confocal imaging of PAR and SUMO-2/3 in U2OS cells upon treatment with MG132 (20 μM, 4h). Plots at the bottom right represent fluorescence intensities of PAR and SUMO-2/3 from condensates indicated by the dashed line. (d) Immunoblots of K6-, K48-, and K63-linked ubiquitin chains in U2OS cells following 4-h incubation in the absence and presence of MG132 (20 μM), BTZ (10 μM), or CFZ (10 μM). (e) Confocal microscopy of PAR and ubiquitin chains in U2OS cells treated with 20 μM MG132 for 4 h. Plots at the bottom right show fluorescence signals of ubiquitin and PAR from condensates indicated by the dashed line. (f) Confocal microscopic analysis of PAR and K63-linked ubiquitin chains in U2OS cells treated for 4 h with DMSO or 20 μM MG132. Scale bars, 10 μm.

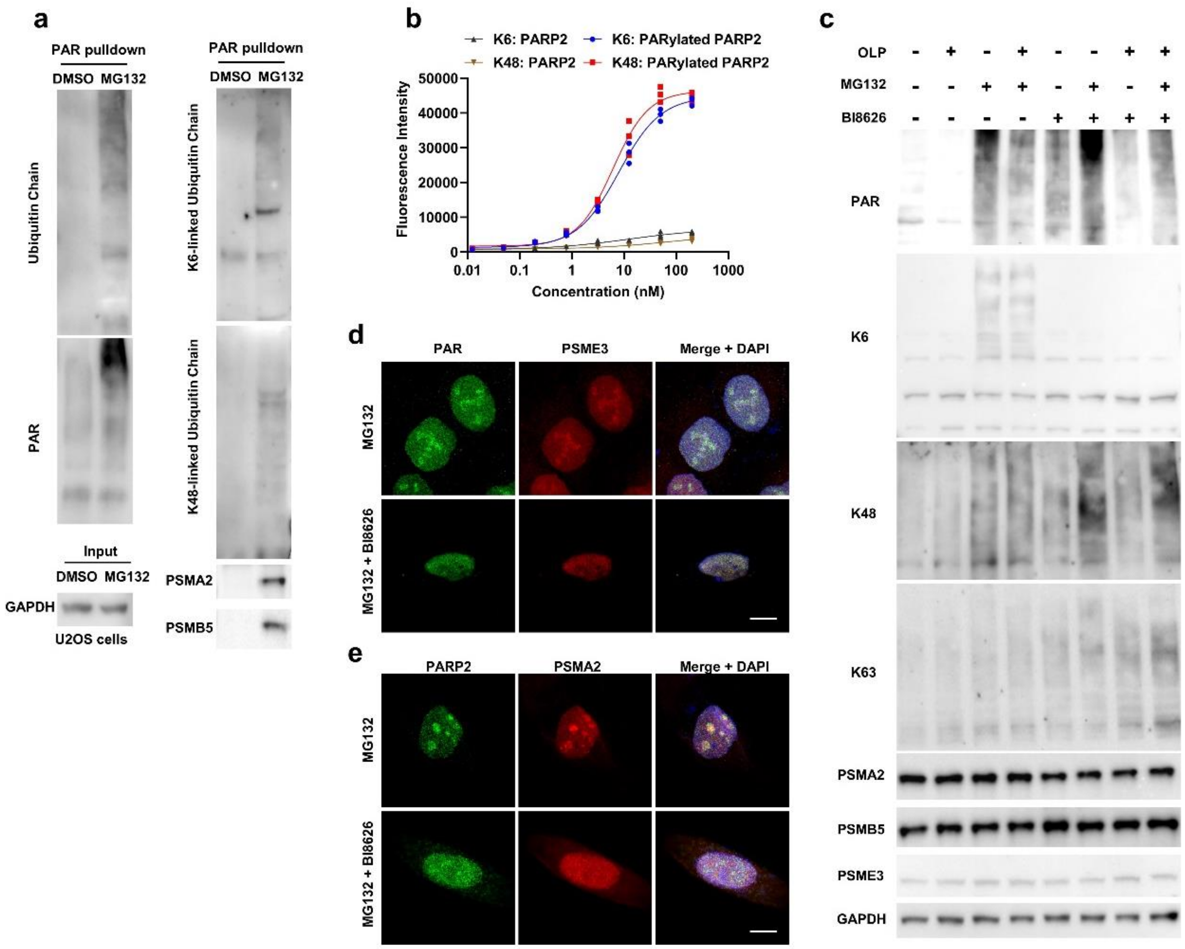


Figure S5. PAR binds to ubiquitin chains and requires K6-linked ubiquitin chains for formation of condensates. (a) Immunoblots of PAR pulldown samples from lysates of U2OS cells treated with DMSO or 20 μM MG132 for 4 h. PAR was precipitated by an anti-PAR monoclonal antibody, followed by immunoblot analysis. (b) Sandwich ELISA analysis of binding of purified PARP2 and PARylated PARP2 to ubiquitin chains. Tetra-K6/K48 ubiquitin chains and anti-PARP2 antibody were utilized as capture and detection reagents, respectively. (c) Immunoblot analysis of U2OS cells without and with treatment by OLP, MG132, and/or BI8626. (d) Confocal imaging of PAR and PSME3 in U2OS cells without and with 4-h treatment by 10 μM BI8626 in the presence of 20 μM MG132. (e) Confocal microscopy of PARP2 and PSMA2 in MG132-treated U2OS cells without and with BI8626 (10 μM). Scale bars, 10 μm.

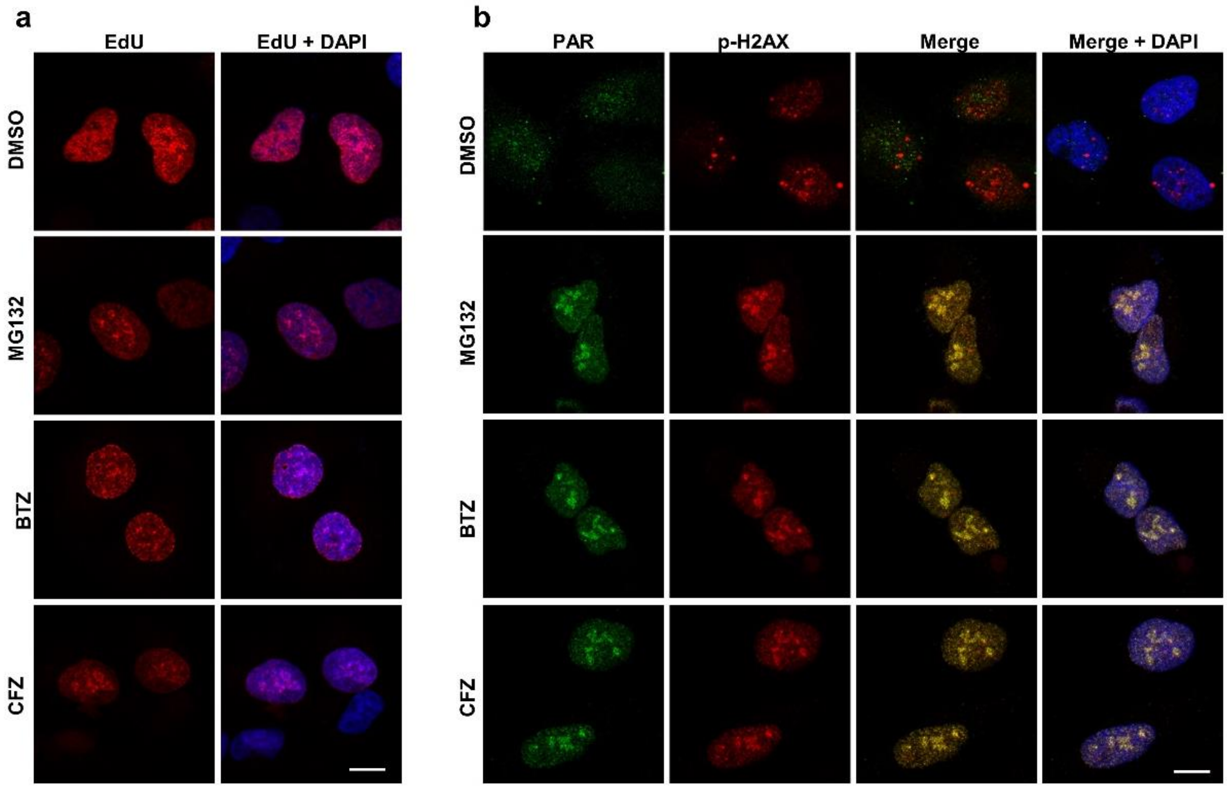


Figure S6. Co-localization of PAR condensates with replication-associated DNA damage sites. (a) Confocal microscopy of EdU incorporation in U2OS cells treated for 4 h by DMSO, 20 μM MG132, 10 μM BTZ, or 10 μM CFZ. (b) Confocal imaging of PAR and p-H2AX in U2OS cells following 4-h treatment with DMSO, MG132 (20 μM), BTZ (10 μM), or CFZ (10 μM). Scale bars, 10 μm.

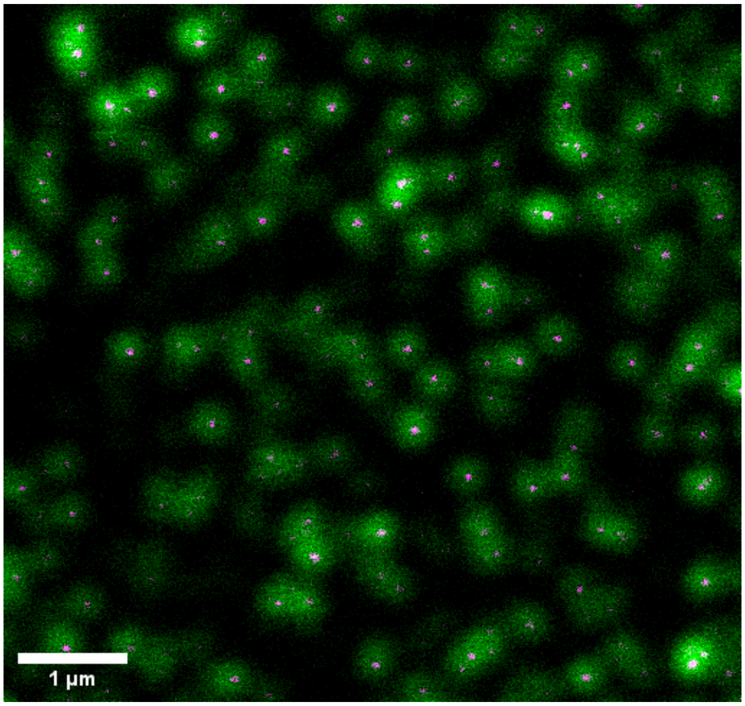


Figure S7. An image was collected to ensure optimal spatial overlap of the excitation and depletion beams using 20 nm beads. Green (confocal image). Purple (STED image).
